# Lower-limb coordination changes following a six-week training intervention that elicited enhancements to maximum velocity sprint performance

**DOI:** 10.1101/2024.06.07.597947

**Authors:** Daniel Lenthall, Adam Brazil, Adrián Castaño-Zambudio, Harry Lightfoot, Jurdan Mendiguchia, Pedro Jiménez-Reyes, Steffi L. Colyer

## Abstract

Alterations to intra- and inter-limb coordination with improved maximal velocity performance remain largely unexplored. This study quantified within-day variability in lower-limb segmental coordination profiles during maximal velocity sprinting and investigated the modifications to coordination strategies in 15 recreationally active males following a six-week period comprised of a multimodal training programme (intervention group (INT); n = 7) or continued participation in sports (control group; n = 8). The INT demonstrated a large decline (effect size = -1.54) in within-day coordination profile variability, suggesting potential skill development. Thigh-thigh coordination modifications for the INT were characterised by an earlier onset of trail thigh reversal in early swing (26 vs. 28% stride) and lead thigh reversal in late swing (76 vs. 79% stride), rather than increases in overall time spent in anti-phase. Moreover, an increase in backwards thigh-dominant, thigh-shank (effect size, 95% CIs: 0.75, 0.17 to 1.33) and shank-dominant, shank-foot (0.76, -0.17 to 1.68) rotations during late swing likely facilitated more aggressive acceleration of the foot prior to touchdown, contributing to reduced touchdown distance and more favourable lower-limb configuration at initial ground contact. These novel findings provide empirical support for the role of longitudinal coordination modifications in improving maximal velocity performance.

**SUMMARY STATEMENT:** Coordination during the swing phase was more modifiable than during stance, with earlier reversal of antiphase thigh-thigh and backwards thigh-shank and shank-foot rotations in late swing observed with improved performance.

## INTRODUCTION

Sprinting is an integral component of many sports and involves a phase of acceleration followed by maximal velocity running, where the velocity achieved is a primary determinant of an athlete’s 100-m sprint time (Slawinski et al., 2017). Whilst the stance limb plays a critical role in the generation of ground reaction force during ground contact, which ultimately dictates increases in (or maintenance of) velocity, the generation of force is also dependent on the athlete’s motion prior to ground contact and at touchdown (Clark et al., 2020). Hence, insights to the movement pattern throughout a stride are warranted to enhance understanding of the task-specific swing phase and its contribution to sprinting performance.

Existing literature has extensively quantified isolated joint and segment kinematics during maximal velocity (Belli et al., 2002; Ito et al., 2008; Miyashiro et al., 2019; Yada et al., 2011; Toyoshima & Sakurai, 2016; Clark et al., 2020), providing valuable insight into sprint running mechanics. For example, Clark et al. (2020) demonstrated a strong positive linear relationship between average thigh angular velocity across the entire stride cycle and ground reaction force production during subsequent stance, indicating the importance of thigh angular velocity for achieving increased maximal velocity. Additionally, the ‘front-side’ mechanics framework proposed by Mann and Murphy (2015) suggests that swing phase thigh rotations are crucial to maximal sprint velocity, with recent predictive simulations supporting the importance of reducing thigh extension in early swing (Haralabidis et al., 2022). However, an evaluation of how functionally linked system components move relative to each other (i.e. their coordination) to satisfy the demands of a given task is also crucial to further understanding of sprint mechanics and the technique developments required to enhance performance (Bezodis et al., 2019).

During accelerative (Bayne, Bezodis & Donaldson, 2020; Donaldson, Bezodis & Bayne, 2022; Donaldson, Bezodis & Bayne, 2024), maximal velocity (Clark et al., 2020) and incline (Okudaira et al., 2021) sprinting, inter-limb thigh-thigh coordination is predominantly anti-phase (opposing rotation). In fact, vector coding analysis of thigh-thigh coupling during accelerative sprinting has found that elite sprinters spend more relative time in anti-phase motion compared with their sub-elite counterparts (Bayne et al., 2020), and that total anti-phase motion decreases progressively from step one to three (Donaldson et al., 2022). However, maximum velocity sprinting has unique task demands compared to acceleration, with the requirement for higher force production in a shorter period of time from a more upright running posture compared with acceleration (Rabita et al., 2015; Von Lieres Und Wilkau et al., 2020). Currently, vector coding analyses of inter- and intra-limb segmental coordination at maximal velocity is missing from the literature, with the study by Gittoes and Wilson (2010) investigating intra-limb joint coordination using continuous relative phase (CRP) with specific focus on CRP variability throughout the stride cycle. Furthermore, previous coordination research has predominantly focused on cross-sectional analysis of sprinting (Donaldson, et al., 2022, 2024; Okudaira et al., 2021; Gittoes & Wilson, 2010), often grouping based on performance level (Bayne et al., 2020) or discipline (Bezodis et al., 2019). Given the self-organising nature of coordination (Newell, 1985), cross-sectional analyses may not represent within-individual changes as individuals within the same group can develop unique coordination strategies to achieve similar task goals (Needham et al., 2014, 2020; Donaldson, et al., 2022, 2024). Therefore, a gap exists in the literature applying longitudinal research design to understand how changes in coordination may facilitate performance improvements.

Inherently connected with coordination analyses is the understanding of variability. Within the context of sports performance, elite athletes exhibit variability in movement patterns even after years of practice (Bartlett et al., 2007; Bradshaw, Maulder, & Keogh, 2007; Davids et al., 2003), indicating that there is an opportunity for inherent biological variability to be promoted in a functional manner within elite sport performance (Robins, et al., 2006). Based on Newell’s stages of learning (Newell, 1985), a non-linear, U-shaped model may exist for the magnitude of variability as skill acquisition progresses, which has been experimentally supported in elite triple jumping (Wilson et al., 2008). The non-linear relationship between task expertise and variability suggests how greater variability might initially be utilised to explore patterns of coordination, before reducing to decrease motor system complexity in task accomplishment (Newell & Vaillancourt, 2001), and later increasing to provide adaptability in responding to perturbations (Wilson et al., 2008). Within maximal velocity sprinting, coordination variability has only been assessed using CRP methods to intra-limb joint couples, indicating touchdown produced greater within-athlete variability than toe-off and that swing produced a less stable coordinative state than ground contact. Thus, at maximal velocity, the swing phase appears to be more modifiable in the pursuit of increased performance.

Currently, our understanding of coordination variability is mostly limited to well-trained populations (Bradshaw, Maulder, & Keogh, 2007; Donaldson et al., 2022; Gittoes & Wilson, 2010; Trezise, Bartlett & Bussey, 2011), and investigation of how variability changes within amateur athletes undertaking a specific sprinting intervention is lacking. Quantifying coordination variability may also be used to define meaningful longitudinal changes, where observed changes outside of this variability could be considered ‘true’ modifications to coordination. Attempts have been made to assess within-day variability in coordination during submaximal treadmill running (Stock et al., 2017) and the velocity ellipse area method (Stock et al., 2018). However, applying both approaches demonstrated poor repeatability of variability during a cutting task, providing potential scope for alternative approaches to assess variability of coordination profiles during maximal velocity sprinting.

Outcomes of a recent six-week multimodal intervention study conducted by our group (Mendiguchia et al., 2022) suggest that combining lumbopelvic control exercises with a sprint technique training program improves performance at maximal velocity. The study observed faster 25-35 m split times and modifications to the sagittal plane kinematics of the lower limb according to the front-side mechanics principles, including higher vertical knee position and thigh angular velocity, and shorter touchdown distance and ground contact duration. However, any modification to coordinative strategies in response to this training intervention are yet to be characterised and would provide novel information to the sprint biomechanics field. The aims of this study were therefore to quantify within-day variation in segmental coordination profiles and utilise these quantities to identify differences in coordination strategies with improved sprint performance. Based on previous research, we hypothesised that the swing phase coordination profiles would be subject to greater modification compared to the ground contact phase and that the multimodal training intervention would increase total thigh-thigh antiphase motion across a stride at maximal velocity.

## RESULTS

The intervention (INT) group improved their maximum velocity from 8.49 ± 0.46 to 8.93 ± 0.53 (effect size = 0.89, 95% CIs = 0 to 1.78) across the training period (PRE to POST), whilst there was a trivial change in maximal velocity exhibited by the control (CON) group (8.91 ± 0.46 to 8.84 ± 0.55). The INT group exhibited decreased mean intra-day coupling angle difference (CAD; effect size = -1.54, 95% CIs = -2.53 to -0.55) at POST (2.65 ± 0.53%) compared with the PRE (4.41 ± 1.90%), whereas changes were small (effect size = -0.11, 95% CIs = -0.68 to 0.46) for the CON group (4.98 ± 1.48% and 4.86 ± 0.73% for PRE and POST, respectively). Meaningful PRE-POST coordination changes (i.e. those that exceeded the mean intra-day variability) were displayed across all segment couples for both groups, with swing phases displaying greater differences in coordination profiles compared with ground contact phases (effect size = 4.63 and 4.03 for the INT and CON group, respectively; Table 1).

**Table 1.** Intra-day and PRE-POST coupling angle differences (CAD, %) during a complete stride at maximal velocity for the intervention and control group.z.

|  | Intervention |  |  |  |  |  | Control |  |  |  |  |  |
| --- | --- | --- | --- | --- | --- | --- | --- | --- | --- | --- | --- | --- |
|  | Intra-day CAD (%) |  |  | PRE-POST CAD (%) |  |  | Intra-day CAD (%) |  |  | PRE-POST CAD (%) |  |  |
|  | PRE | POST | Mean | Contact | Swing | Stride | PRE | POST | Mean | Contact | Swing | Stride |
| **Thigh-Thigh** | 6.06 ± 3.94 | 2.85 ± 0.79 | 4.46 ± 2.27 | 0.78 ± 0.27 | 6.96 ± 2.61 | 7.75 ± 2.45* | 6.44 ± 4.98 | 5.12 ± 2.52 | 5.79 ± 0.93 | 0.98 ± 0.51 | 5.85 ± 2.72 | 6.83 ± 3.15* |
| **Thigh-Shank** | 2.90 ± 0.94 | 2.05 ± 0.74 | 2.56 ± 0.47 | 1.10 ± 0.79 | 4.91 ± 2.92 | 6.01 ± 2.97* | 3.49 ± 1.61 | 4.03 ± 1.58 | 3.76 ± 0.38 | 0.71 ± 0.43 | 4.80 ± 1.56 | 5.52 ± 1.64* |
| **Shank-Foot** | 4.28 ± 2.30 | 3.04 ± 0.97 | 3.66 ± 0.88 | 1.07 ± 0.52 | 6.64 ± 3.68 | 7.70 ± 3.79* | 5.02 ± 3.09 | 5.42 ± 1.91 | 5.22 ± 0.27 | 0.64 ± 0.38 | 6.32 ± 1.78 | 6.96 ± 1.72* |
| **Mean across all couples** | 4.41 ± 1.58 | 2.65 ± 0.53 | 3.53 ± 1.00 | 0.98 ± 0.18 | 6.17 ± 1.10 | 7.15 ± 0.99* | 4.98 ± 1.48 | 4.86 ± 0.73 | 4.92 ± 1.04 | 0.78 ± 0.18 | 5.66 ± 0.78 | 6.44 ± 0.80* |

\*denotes meaningful PRE-POST coordination changes (i.e. those that exceeded the intra-day variability).

For thigh-thigh coordination (Figure 1), the INT group reduced the proportion of stride spent in anti-phase (-+) ipsilateral thigh dominance (-0.57, -1.32 to 0.19, small; ES, CIs, ES classification) and increased anti-phase (-+) contralateral thigh dominance (0.57, -0.19 to 1.32, small) during ipsilateral ground contact. An earlier onset of contralateral (lead) thigh (anti-clockwise to clockwise rotation) and ipsilateral (trail) thigh reversals (26 vs. 28% stride, 29 vs. 30% stride, respectively) during ipsilateral early swing resulted in reduced in-phase (--) contralateral thigh dominance (-0.38, -0.93 to 0.17, small) coordination. During contralateral ground contact, the CON group elicited changes in anti-phase (+-) coordination, with decreased ipsilateral thigh dominance (-1.08, -2.28 to 0.11, moderate) and increased contralateral thigh dominance (1.08, -0.11 to 2.28, moderate). During contralateral early swing, the INT group exhibited earlier transitions from anti-phase (+-) to in-phase (--) coordination (76 vs. 79% stride) and in-phase (--) to anti-phase (-+) coordination (80 vs. 79% stride), with reduced anti-phase (+-) ipsilateral thigh dominance (-1.24, -2.20 to -0.27, large). The INT group also reduced in-phase (++) contralateral thigh dominance (-0.30, -0.67 to 0.08, small) in contralateral early swing, although closer inspection revealed this small effect is due to one individual exhibiting a large change. In line with the INT group, the CON group exhibited reduced anti-phase (+-) contralateral thigh dominance (-0.77, -1.56 to 0.03, moderate) coordination after contralateral toe-off. However, in contrast to the INT group, the CON group increased anti-phase (-+) coordination (Figure 1), highlighted by increased anti-phase (-+) contralateral thigh dominance (1.02, 0.00 to 2.05, moderate) coordination throughout contralateral early swing.

**Figure 1.**
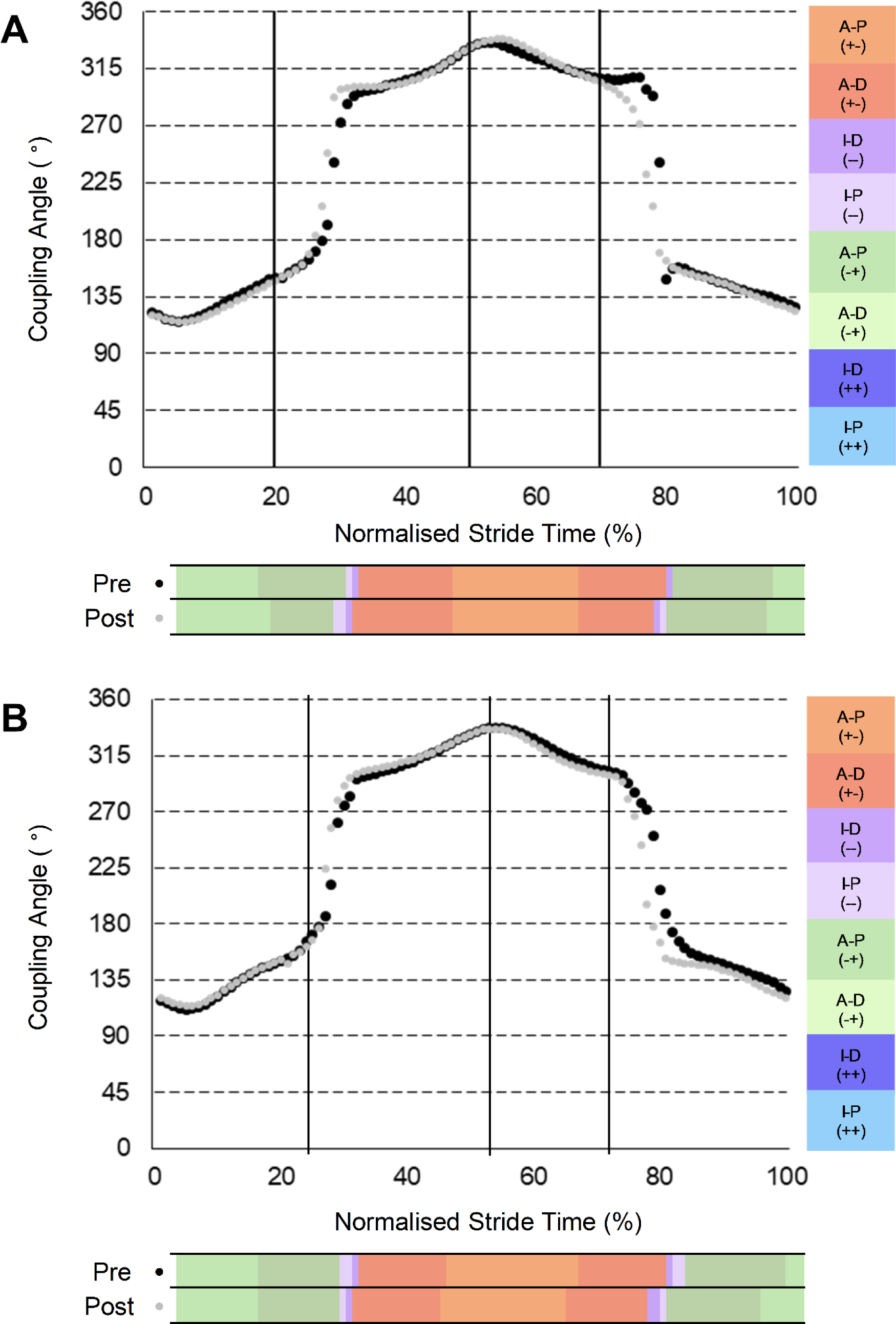
Group-mean PRE and POST thigh-thigh coupling angles normalised to a stride of maximal velocity sprinting. (A) Intervention group (n = 7). (B) Control group (n = 8). Black dots denote PRE; grey dots denote POST.

For thigh-shank coordination (Figure 2), during ipsilateral early swing, both groups showed an earlier onset of anti-clockwise thigh rotation, resulting in slightly earlier (29 vs. 30% stride) transitions away from in-phase (--) shank dominant coordination (INT: -0.71, -1.45 to 0.03, moderate; CON: -0.30, -0.81 to 0.20, small). The INT group subsequently increased anti-phase (+-) thigh dominance (1.31, 0.32 to 2.31, large) coordination. In contrast, the CON group increased in-phase (++) thigh dominance (0.58, -0.01 to 1.16, small) coordination. During contralateral ground contact, earlier reversal of shank rotation (52 vs. 54% stride) for the INT group served to increase in-phase (++) thigh dominance (0.38, 0.12 to 0.64, small) and reduce anti-phase (+-) thigh dominance (-0.88, -1.69 to -0.06, moderate) coordination. An earlier onset of clockwise thigh rotation after contralateral toe-off (76 vs. 79% stride) for the INT group reduced in-phase (++) shank dominant (-1.19, -2.21 to -0.18, moderate) and anti-phase (-+) shank dominant (-0.62, -1.35 to 0.12, moderate), but increased anti-phase (-+) thigh dominant (0.75, 0.17 to 1.33, moderate) coordination. The CON group also transitioned earlier (77 vs. 79% stride) away from in-phase (++) shank dominant (-0.91, -1.73 to -0.09, moderate) coordination (Figure 2). However, in contrast to the INT group, the CON group displayed no change in anti-phase (-+) but increased in-phase (--) shank dominant (1.07, 0.54 to 1.61, moderate) coordination.

**Figure 2.**
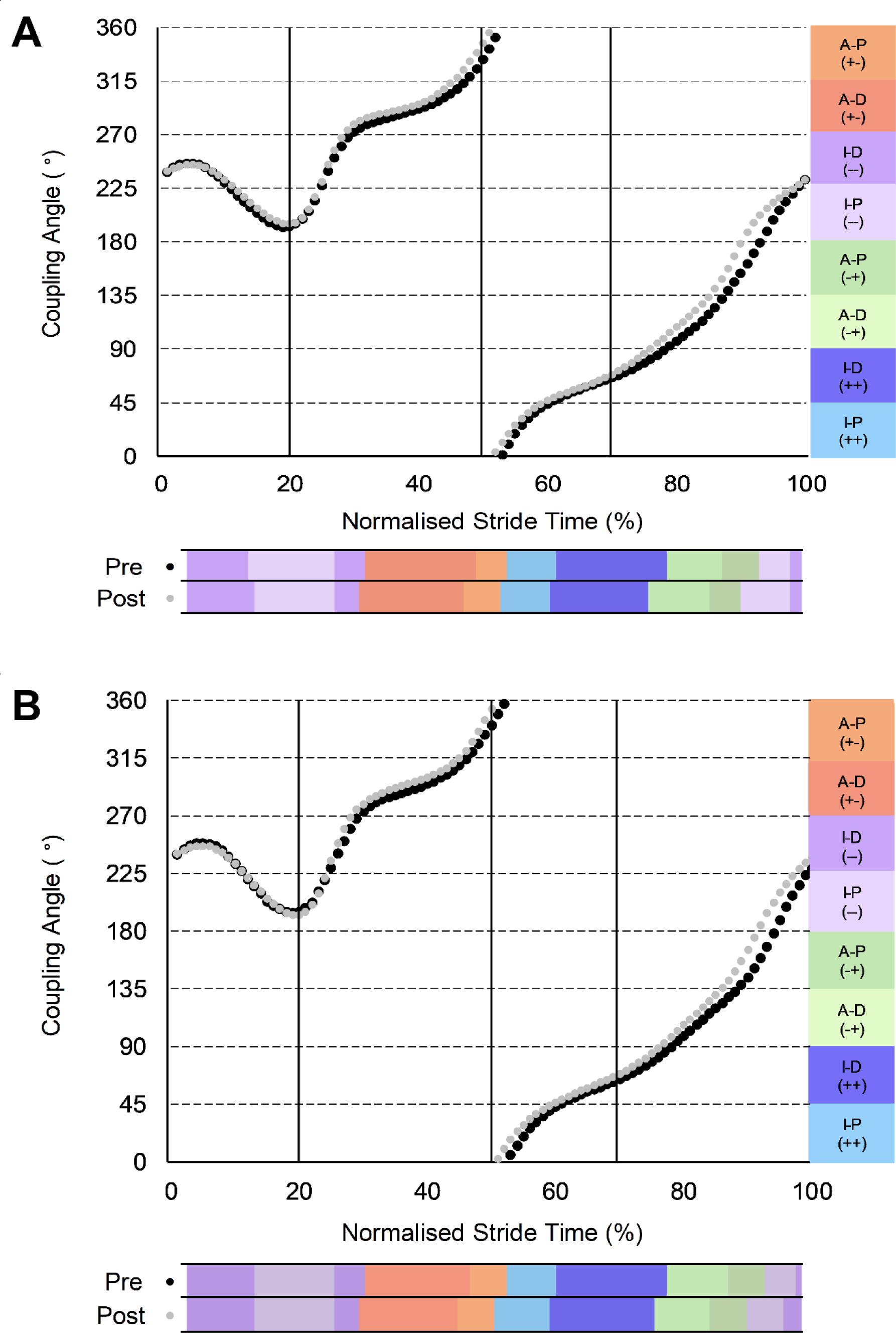
Group-mean PRE and POST thigh-shank coupling angles normalised to a stride of maximal velocity sprinting. (A) Intervention group (n = 7). (B) Control group (n = 8). Black dots denote PRE; grey dots denote POST.

For shank-foot coordination (Figure 3), during ipsilateral early swing, the CON group increased in-phase (++) foot dominance (0.58, -0.01 to 1.16, small) and anti-phase (-+) shank dominance (0.40, -0.10 to 0.89, small) coordination. Moreover, earlier shank reversal (clockwise to anti-clockwise rotation) in initial contralateral ground contact l (49 vs 48% stride) for both groups resulted in reduced anti-phase (-+) shank dominance (INT: -0.45, -1.05 to 0.16, small; CON: -0.58, -1.16 to 0.01, small) for both groups, reduced anti-phase (-+) foot dominance (-0.43, -0.86 to 0.01, small) for the CON group, and increased in-phase (++) foot dominance (0.96, 0.12 to 1.81, moderate) and reduced in-phase (--) shank dominance (-0.73, - 1.45 to -0.02, moderate) for the INT group. Earlier onset of foot (88 vs. 91% stride) and shank (91 vs. 94% stride) clockwise rotations during contralateral early swing for the INT group resulted in an earlier transition away from in-phase (++) foot dominance (-1.52, -3.02 to -0.01, large) through to longer duration in-phase (++) shank dominance (0.67, -0.03 to 1.36, moderate), anti-phase (+-) shank dominance (0.34, 0.03 to 0.64, small) and in-phase (--) shank dominance (0.76, -0.17 to 1.68, moderate) coordination. In contrast, the CON group displayed earlier transition (92 vs. 95% stride) from anti-phase (+-) to in-phase (--) coordination (Figure 3), emerging as reduced anti-phase (+-) shank dominance (-0.29, -0.68 to 0.10, small) and increased in-phase (--) foot dominance (1.12, 0.18 to 2.07, moderate) coordination.

**Figure 3.**
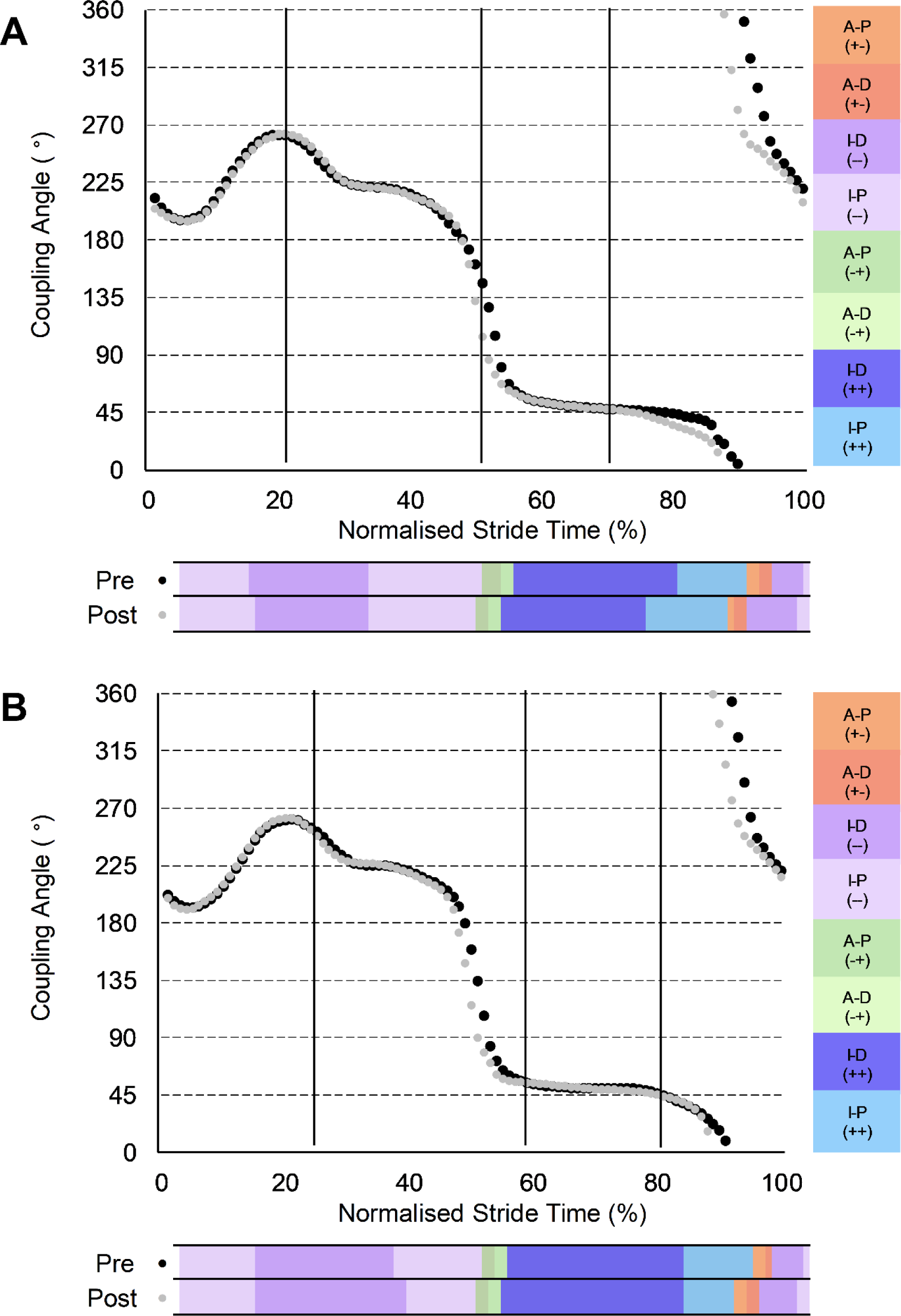
Group-mean PRE and POST thigh-shank coupling angles normalised to a stride of maximal velocity sprinting. (A) Intervention group (n = 7). (B) Control group (n = 8). Black dots denote PRE; grey dots denote POST.

## DISCUSSION

The aims of this study were to quantify within-day variation in lower-limb segment coordination profiles at maximal velocity and to identify differences in coordination strategies with improved sprint performance following a six-week multimodal training program. A meaningful decrease (effect size = -1.54) in coordination profile variability, analysed by using coordination binning (Needham et al., 2020) and the CAD method (Brazil et al. 2020), was observed in the INT group post-intervention, while a small difference (effect size = 0.34) was observed in the CON group, supporting that a reduction in coordination variability and a more fixed sprinting pattern can be associated with skill acquisition. Additionally, this study observed that during the swing phase participants exhibited increased PRE-POST coordination profile changes relative to GCT across all couples for the INT group (mean ± SD; swing: 6.64 ± 2.46%, GCT: 1.29 ± 0.51%) and the CON group (swing: 5.84 ± 1.62%, GCT: 1.41 ± 0.69%), implying that changes to coordination strategies are primarily afforded during the swing phase of a maximal velocity stride. From an injury risk perspective, changes in coordination in late swing of a maximum velocity stride may have important implications as this is when the hamstrings are most susceptible to injury from excessive lengthening (Chumanov et al., 2011). This study also observed, coinciding with improved performance, no difference in total anti-phase thigh-thigh motion across a stride at maximal velocity but an evident occurrence of earlier thigh rotation reversals, adoption of a more ‘front-side mechanics’ movement pattern and more aggressive retraction of the lower-limb in the late swing.

For the INT group, intra-day variability in coordination profiles decreased POST relative to PRE, indicating a reduction in coordination profile variability. This reduction was concomitant with improved sprint performance for the INT group, which is consistent with Newell’s stages of learning (Newell, 1985) and research indicating that as athletes progress in expertise from a novice level, movement execution is more consistent (Huys, Daffertshofer & Beek, 2003; Wilson et al., 2008; Yang & Scholz, 2005) and may require less conscious attention demand (Beilock et al., 2004). However, the previously established U-shaped relationship between task expertise and coordination variability (Wilson et al., 2008) must be considered, and longitudinal studies are necessary to gain insights on the functionality of coordination variability as expertise progresses at the highest performance levels.

From a constraints-based approach (Newell, 1986), this study indicates that maximal velocity sprinting has relatively strong task constraints which yield broadly similar coordination patterns across a sample of amateur male athletes. However, task constraints do not appear stable across all phases of the stride. High consistency across all segment couplings was observed during ground contact but within-individual variation in the timing of transitions between coordination patterns presented during swing. Maximal velocity sprinting may therefore be constrained to an invariant coordination strategy during the closed-chain ground contact phase, indicative of self-organisation towards task-specific coordinative structures (Newell, 1986)-the task being to first attenuate a rapidly occurring impact force at touchdown and subsequently decelerate the centre of mass as quickly as possible to maximise propulsive force generation (Gittoes & Wilson, 2010). Greater degrees of freedom afforded to athletes to flexibly modify coordination strategies during swing suggests interventions aimed at enhancing maximal velocity performance may wish to target modifying movement execution during the swing phase with the view to contacting the ground in a more favourable position for force transmission.

Thigh-thigh coordination was primarily anti-phase across a stride at maximal velocity (Figure 1), supporting the oscillatory thigh motion reported by Clark et al. (2020). The high frequency of anti-phase coordination also aligns with the observations of Bayne et al. (2020) and Donaldson, et al. (2022, 2024) in accelerative sprinting, and Okudaira et al. (2021) during incline sprinting. However, in line with accelerative sprinting (Bayne et al., 2020; Donaldson et al., 2022), no participant exhibited a perfect scissor (continuous anti-phase) motion. Yet, contrary to accelerative sprinting where oscillatory thigh motion is asymmetric and trail leg dominant (Donaldson et al., 2022), thigh motion during maximal velocity sprinting appears symmetric (Figure 1). This likely reflects specific task constraints between acceleration and maximum velocity sprinting, including the transition from blocks to overground sprinting, the lower absolute velocity, and greater accelerative demand. Increased anti-clockwise contralateral thigh (swing) rotation relative to clockwise ipsilateral thigh rotation in ipsilateral ground contact for the INT group may have served to improve trail thigh recovery. An earlier initiation of ipsilateral thigh (trail thigh now) reversal (clockwise to anti-clockwise rotation) in the INT group during swing is aligned with a more rapid recovery of the swing limb, reduced backside mechanics and increased mean thigh retraction velocity (Mendiguchia et al., 2022). Further, earlier trail thigh pull suggests increased thigh retraction velocity which has been associated with faster running speeds at maximal velocity (Clark et al., 2020). Nagahara et al. (2017) have shown that increments in velocity are associated with increased negative work of the hip during early swing and the knee during terminal swing, which may be underlying causes of the kinematic modifications observed in the current study, although more research is required to fully elucidate this.

During contralateral early swing, the INT group displayed decreased time in anti-clockwise ipsilateral (lead) thigh rotation, indicating earlier attainment of maximal vertical position of the lead knee, a more ‘front-side’ profile and earlier onset of lead limb reversal. Earlier retraction of the lead limb in late swing in line with faster average thigh angular velocity over the stride (as shown previously within these participants, Mendiguchia et al., 2022) suggests earlier initiation of accelerating the lead thigh backwards into touchdown. Such a change is associated with “punching” the swing leg into ground contact, facilitating increased thigh angular velocities and increased force production in subsequent ground contact (Clark et al. 2020). Considering Morin et al. (2015) reported that increased activation of the hamstring muscles late in swing augments force production upon ground contact during acceleration, the increased average thigh angular velocity across the stride observed previously (Mendiguchia et al., 2022) could be attributed to increased activation of the hamstring muscles late in swing as a result of earlier retraction of the lead limb. Additionally, we speculate that this more favourable coordination pattern may be attributable to smoother switching between rectus femoris and biceps femoris muscle activations, as this has previously been correlated with step frequency at maximal velocity (Kakehata et al., 2021). However, considering the multimodal intervention incorporated S&C training for two of the weekly sessions, coordination changes late in swing may stem from the heightened physical strength of the participant’s POST. Therefore, varying strength capabilities could potentially be linked to distinct patterns of thigh-thigh coordination, although further research is needed.

Bayne et al. (2020) reported that the proportion of anti-phase coordination was higher in elite athletes (85.9%) than in sub-elite athletes (76.8%) during accelerative sprinting. However, the current study observed no meaningful PRE-POST change in total anti-phase thigh-thigh motion across the stride for the INT group (0.192, -0.688 to 1.071, trivial). Rather, the primary PRE-POST difference was one of timing; at POST, participants displayed an earlier onset of ipsilateral and contralateral thigh reversals both in ipsilateral early swing and contralateral early swing, reflecting earlier trail thigh pull and lead thigh retraction, respectively. Importantly, in agreement with the current study, a recent study demonstrated that higher-performing sprinters exhibit earlier swing thigh rotation reversal but no difference in total thigh-thigh anti-phase motion in steps 2-4 of acceleration (Donaldson et al., 2024). Early swing thigh rotation reversal therefore appears key in both accelerative and maximum velocity sprinting.

For thigh-shank coordination, the INT group reduced clockwise and increased anti-clockwise thigh rotation during ipsilateral early swing, reflecting reduced backside mechanics, which has recently been shown via predictive simulations to be associated with improved sprint acceleration performance (Haralabidis et al., 2022). During contralateral early swing (lead thigh retraction after maximum thigh separation) an earlier onset of clockwise thigh rotation was observed for the INT group (Figure 2). These findings imply increased time to accelerate the foot down and back before touchdown, offering a possible explanation for the previously observed reduction of touchdown distance in the INT group (Mendiguchia et al., 2022). Furthermore, the earlier shank and foot reversals for the INT group (Figure 3) suggest relatively longer clockwise shank-foot rotation in preparation for touchdown, likely reflecting the early lead limb retraction during this phase. Hence, the foot moved further backwards before touchdown, resulting in a reduced touchdown distance (Mendiguchia et al., 2022) and potentially improving stiffness of the ankle during stance, minimising positive rotation of the foot and aiding force transmission. A smaller touchdown distance is associated with lower magnitudes of relative braking impulse during the early part of ground contact in the mid-acceleration phase (Hunter, Marshall & McNair, 2005) and could intuitively also be favourable at maximal velocity, where attenuating braking forces is a crucial performance determinant (Colyer et al., 2018). Intuitively, the alterations to the shank-foot coupling could increase the stiffness of the ankle during the subsequent ground contact, which has been associated with maximal sprint velocity improvements (Nagahara & Zushi, 2017), and would likely reduce the ground contact time, which was observed previously (Mendiguchia et al., 2022).

The current study is the first to adopt a longitudinal approach to assess coordination strategy alterations at maximal velocity and has provided new insights into coordination changes associated with performance improvement. Nevertheless, the limited sample size (n=15) of amateur athletes restricts the extrapolation of findings to elite populations. Indeed, a ‘ceiling effect’ may exist for elite athletes already operating near their peak performance (Newell, 1985). Future research should therefore investigate coordination modifications in elite sprinters (and/or players in other sports involving sprinting e.g. rugby and football) following participation in interventions tailored to address specific weaknesses. The use of CAD to assess coordination variability in the current study is novel in approach and offers a holistic way to understand variability in the coordination profile. Whilst coordination variability is typically assessed through circular variability of raw coupling angle data (Needham et al., 2020), or bivariate solutions that can be less prone to statistical artefact (e.g., Stock et al., 2018), the current solution may offer an alternative approach to understanding variability in coordination profiles rather than the coupling angle itself. Whilst granularity from circular statistics might be reduced, the variability measured through CAD aligns with methods used to practically interpret coordination profiles and may offer protection from statistical artefact at short vector lengths by having a wider bandwidth of tolerance.

## CONCLUSION

The current study has quantified within-day variation in intra- and inter-limb coordination and defined coordination changes at maximal velocity following a six-week multimodal intervention program. Inter-limb thigh-thigh and intra-limb thigh-shank and shank-foot coordination appear modifiable, particularly during the swing phase compared to during ground contact. Most notably, the INT group thigh-thigh decreased in-phase (--) coordination during ipsilateral early swing and anti-phase (+-) coordination during contralateral early swing, facilitating earlier lead and trail thigh reversals. However, in contrast to our hypotheses, there was no difference in the proportion of stride in which anti-phase thigh-thigh motion was exhibited. Nevertheless, decreased clockwise in-phase (++) thigh-shank coordination in late swing possibly improved limb recovery through more aggressive retraction of the foot in preparation for touchdown, which likely contributed to reduced touchdown distance. By identifying key coordinative modifications which distinguish faster maximal velocity sprinting times, this study contributes a new conceptual understanding of the role of coordination modifications in improving maximal velocity sprint performance, which can be utilised when developing technical models of sprinting and inform the design of training interventions.

## MATERIALS AND METHODS

### Participants and data collection procedure

A detailed description of the data collection and processing and multimodal training intervention methodologies have been provided previously (Mendiguchia et al., 2022). In summary, a prospective randomised control trial was conducted, with testing sessions before and after six weeks of training. Fifteen amateur male athletes were assigned in a counterbalanced method according to the initial sprint performance into two groups: eight athletes in the CON group and seven in the INT group. At each testing session, participants maximally sprinted twice over 35 m, with a four-minute recovery period between efforts. During these attempts, 3D lower-limb kinematic data of a single full stride (touchdown to touchdown of the same limb) at maximal velocity were collected using 15 Qualisys Oqus cameras. A capture volume of approximately 10 × 1.1 × 1.5 m was calibrated according to the manufacturer’s guidelines. Twenty-four markers were placed bilaterally on the following lower-limb landmarks: posterior superior iliac spine, anterior superior iliac spine, greater trochanter, medial and lateral femoral condyles, medial and lateral malleoli, heel, first and fifth metatarsophalangeal joints, and the hallux. Additionally, rigid clusters of 4 markers were attached to the thigh and shank segments (see Mendiguchia et al., 2022 for full details). During the six-week training period, the CON group continued participation in sports involving sprinting at least three times a week, whereas the INT group partook in a multimodal training program comprised of three weekly sessions integrating coaching, strength and conditioning (S&C), and physical therapy approaches (full detail provided in supplementary material within Mendiguchia et al., 2022).

### Data processing

A seven-segment kinematic model was created in Visual 3D (version 6; C-Motion Inc, Germantown, MD), with each segment’s (pelvis and bilateral thighs, shanks and feet) coordinate system defined as a right-handed orthogonal coordinate system with three axes: X (medio-lateral), Y (anterior-posterior) and Z (longitudinal). Thigh, shank, and foot segment orientations were calculated relative to the global coordinate system using an XYZ Cardan sequence of rotations, with rotations about the X axis analysed subsequently. Touchdown events were computed utilising the method of Handsaker et al. (2016). Within the stride, four key functional phases were identified and used in subsequent analysis: ipsilateral ground contact, ipsilateral early swing, contralateral ground contact, contralateral early swing. The mean times spent in each of these phases were calculated as a percentage of the stride (Figure 4) and data were normalised accordingly as follows: 0-20% (ipsilateral ground contact), 21-50% (ipsilateral early swing), 51-70% (contralateral ground contact), 71-100% (contralateral early swing). Segment rotations were defined as clockwise (-) or anti-clockwise (+) relative to a left-to-right direction of motion. The maximum instantaneous horizontal velocity of the pelvis segment was also extracted as a proxy measure of the maximum centre of mass velocity. All further data processing was performed in MATLAB (v2023a, MathWorks Inc., Natick, MA, USA).

**Figure 4.**
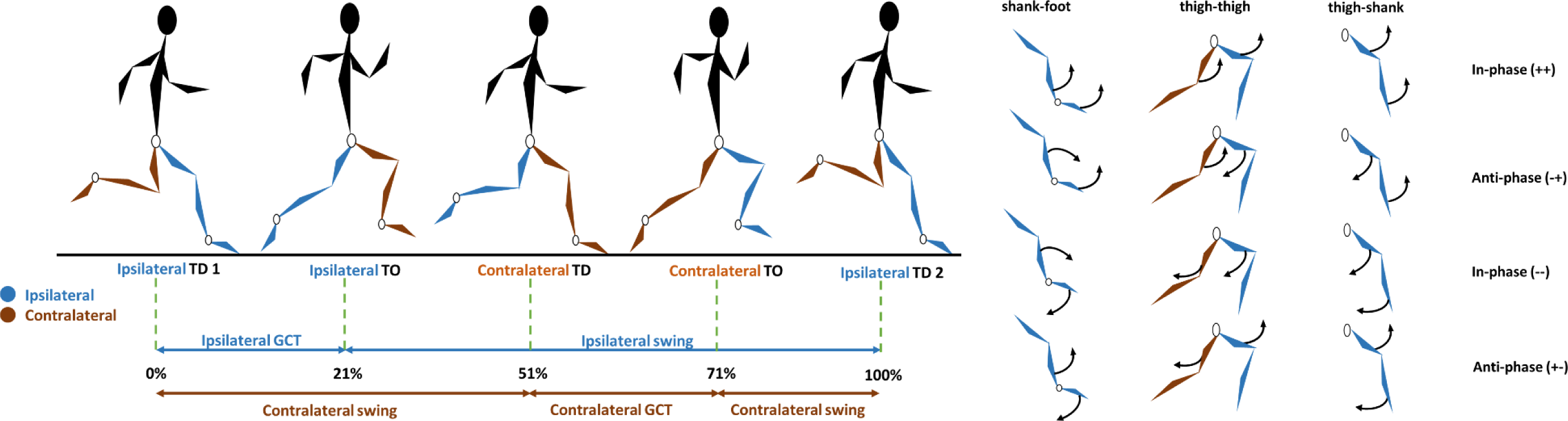
Visual representation of the functional phases within a maximal velocity stride (touchdown to touchdown of the same limb). GC refers to the ground contact phase, TD refers to touchdown, TO refers to toe-off. % values refer to the normalised % of the stride.

Coupling angle mapping was used to profile individual coordination strategies throughout the stride (Needham et al., 2014). Inter-limb thigh-thigh and intra-limb thigh-shank and shank-foot segment couplings were described as proximal-distal. For the thigh-thigh couple, the ipsilateral and contralateral thighs were defined as proximal and distal, respectively. Segment angles were temporally normalised to 101 data points across the stride and vector coding was applied to calculate continuous coupling angles for all segment couples (Needham et al., 2014; Silvernail et al., 2018). Each coupling angle was calculated as the angle of the resultant vector between two consecutive time points on the angle-angle plot, relative to the right horizontal, expressed as an angle between 0° and 360° (Chang, Van Emmerik & Hamill, 2008). To quantify the frequency of coordination patterns, coupling angle data were then classified into one of eight distinct coordination patterns (bins) according to the relationship between segment rotations (in-phase or anti-phase), the direction of rotation (clockwise or anti-clockwise) and the segmental dominance (proximal or distal) (Needham et al., 2020; Figure 5). Coordination patterns were assigned a specific colour (Figure 5) to aid with visualising PRE-POST differences.

**Figure 5.**
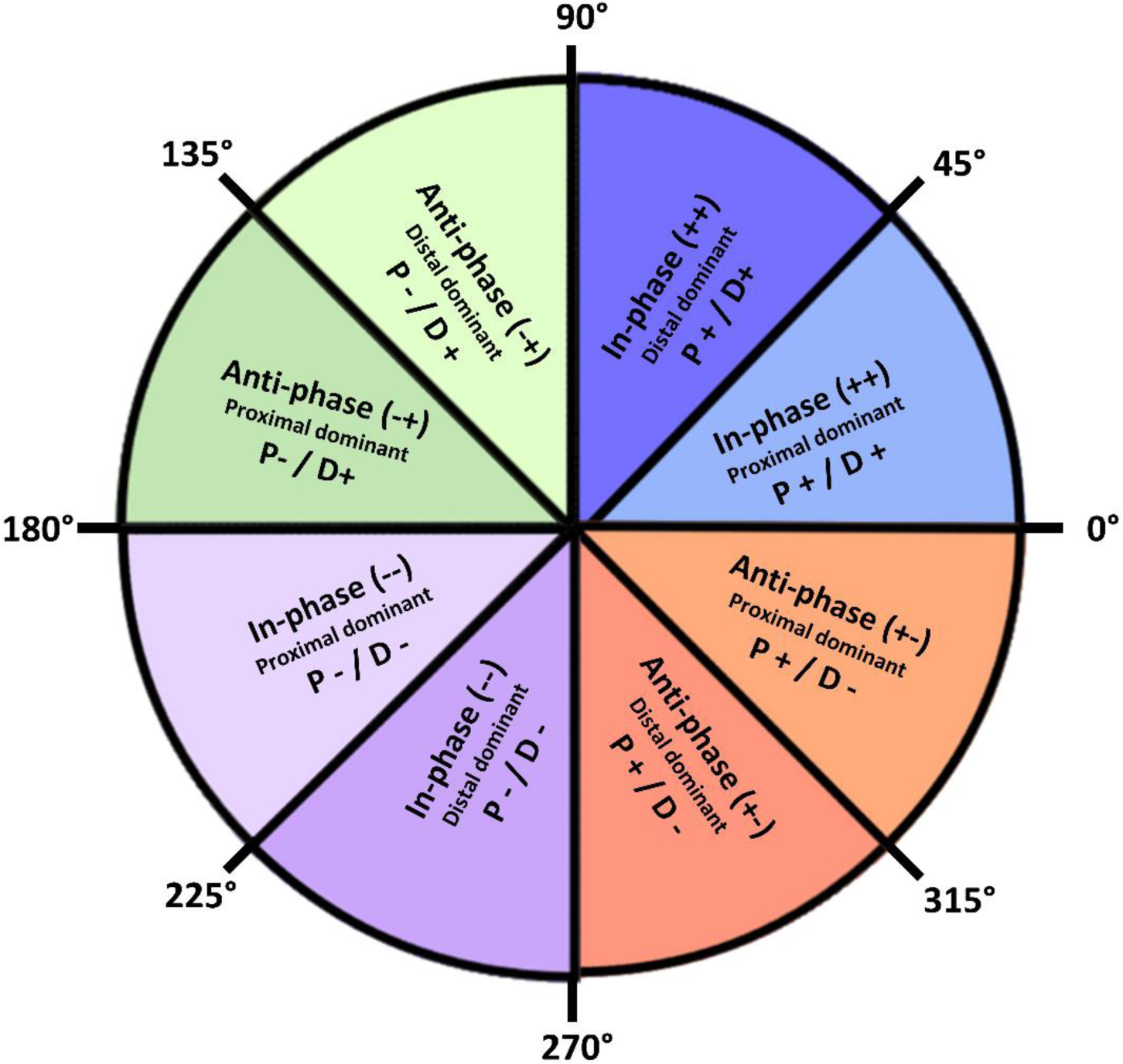
Classification of coordination pattern bins based on the relative motion of each segment (adapted from Needham et al., 2020)

Each athlete’s fastest PRE and POST trial was used for PRE-POST (inter-day) coordination analysis. Both trials were used in the assessment of intra-day coordination profile variability at PRE and POST. To quantify differences in coordination profiles, a coupling angle difference (CAD) score (Brazil et al., 2020) was calculated for each individual and then averaged within each group to quantify PRE-POST coordination changes and intra-day coordination pattern variability. To calculate each CAD, the difference in the coordination bin at each instance of the normalised stride was assigned a score between 0 (same bin) and 4 (opposite bin). The sum of each difference score was represented as a percentage of the maximal possible value (404), with a lower score indicating greater similarity (less variability) in the coordination profile (Bezodis et al., 2019; Brazil et al., 2020). To help make inference on coordination changes, PRE-POST differences were considered meaningful if the group mean PRE-POST CAD was greater than the group mean intra-day CAD. For the presentation of data, all mean coordination profiles were calculated using circular statistics (Needham et al., 2020; Chang, Van Emmerik & Hamill, 2008).

### Statistical Analysis

All within-group, inter-day comparisons were made using group means and standard deviations (SD), and PRE-POST differences were calculated as POST minus PRE. Paired sample t-tests were used to analyse PRE-POST changes in segment ROM and frequency of each coordination bin within the four functional phases. Effect sizes (ES) were calculated using Cohen’s d standardised differences, with mean and pooled SD calculated according to Altman and Gardner (2000), and 95% confidence intervals (CIs) were also computed. ES magnitudes were categorised as small (0.2 ≤ d < 0.6), moderate (0.6 ≤ d < 1.2), large (1.2 ≤ d < 2.0), very large (2.0 ≤ d < 4.0), and extremely large (d ≥ 4.0; Hopkins, 2000). PRE-POST responses were considered practically meaningful when the CIs did not cross 0.2 on the other side of zero to the effect size.

## ACKNOWLEDGEMENTS

The authors sincerely thank the participants for their cooperation and involvement with the study. The authors also thank Professor Neil Bezodis, Dr Helen Bayne and Byron Donaldson for their valuable insights during the initial conceptualization of the analyses.

## AUTHOR CONTRIBUTIONS

Conceptualization and design: SC, JM, PJ; Investigation: SC, AC-Z, Formal Analysis: DL, HL, SC, AB; Project Administration/Supervision: SC, AB, JM, PJ; Writing: DL, AB, AC-Z, HL, JM, PJ, SC.

